# Stratified tissue biofabrication by rotational internal flow layer engineering

**DOI:** 10.1101/2022.12.06.519250

**Authors:** Ian Holland, Wenmiao Shu, Jamie A. Davies

**Affiliations:** Deanery of Biomedical Science and the Centre for Engineering Biology, University of Edinburgh, Edinburgh, United Kingdom; Department of Biomedical Engineering, University of Strathclyde, Glasgow, United Kingdom

## Abstract

The bioassembly of layered tissue that closely mimics human histology presents challenges for tissue engineering. Existing bioprinting technologies lack the resolution and cell densities necessary to form the microscale cell-width layers commonly observed in stratified tissue, particularly when using low-viscosity hydrogels, such as collagen. Here we present rotational internal flow layer engineering (RIFLE), a novel biofabrication technology for assembling tuneable, multi-layered tissue-like structures. Using high-speed rotating tubular moulds, small volumes of cell-laden liquids added to the inner surface were transitioned into thin layers and gelled, progressively building macroscale tubes composed of discrete microscale strata with thicknesses a function of rotational speed. Cell encapsulation enabled the patterning of high-density layers (10^8^ cells/ml) into heterogenous constructs. RIFLE versatility was demonstrated through tunica media assembly, encapsulating human smooth muscle cells in cell-width (12.5μm) collagen layers. This enabling technology has the potential to allow researchers to economically create a range of representative stratified tissue.

## Introduction

Layered tissue is common throughout human anatomy, with notable examples found in cardiac^1^, dermal^2^, intestinal^3^, retinal^4^ and vascular tissue^5^. Such stratification features a range of different cell types and extracellular matrices to create highly specialised, heterogeneous structures that imbue tissues with multiple properties and functions. Layered tissue follows the pattern of many other tissue types where microtissue units combine to form larger macrotissue structures^6^. Tissue engineering technologies, including biofabrication^7,8^, that intend to anatomically and functionally mimic native tissue often make extensive use of cells suspended in liquid-phase hydrogels that can be deposited and subsequently gelled^9^. When using such an approach to form layered tissue, researchers have few technical options. Extrusion-based 3D bioprinting technologies, the mainstays of research and industry^10^, are ill-suited to assembling these hydrogels into planar layered structures. In particular they are unable to attain the required deposition resolution, cell density, repeatability^11,12^ and defined interfaces^13^ needed for assembling the multiple thin continuous layers and hence the hierarchical structures^7,14^ observed in native tissue structures. Such stratified features can often be the width or a cell or smaller^15^. These technologies also impose constraints on the range of compatible hydrogels with printed liquids requiring rapid gelation or support to assume the intended 3D spatial form.

Consequently, a large amount of research effort is devoted to developing printable bioinks^10,16^, the properties of which are frequently aligned to the limitations of the printing technology rather than replicating the form of the target tissue. There remains a need for new biofabrication technologies that are able to form the complex microarchitecture of native tissue^12^.

Recognising the challenges of using existing technologies to manipulate hydrogels into microscale layers, some researchers have attempted to develop improved biofabrication technologies. The sequential, dipping, spraying or direct extrusion of cell-laden hydrogels onto the external surface of cylinders to build up layers is one such strategy^17^. An advantage of this method resides in the formation of a tubular macroarchitecture, observed in many native layered tissues such as vascular, intestinal, tracheal, urethral and biliary tissue^11,18^. Furthermore, cutting open tubular fabricated tissue allows simple conversion to planar structures^19^, as would be needed to mimic dermal and retinal tissue. Rotation of the cylinder at high speeds under motor control^20,21^ enables a level of control over the layer thicknesses that is otherwise determined by the viscosity of the liquid hydrogel and the surface properties of the substrate layer^22^. A limitation of rod-based methods is the requirement to dip or spray cell solutions onto the external surface, necessitating large volumes of cell laden hydrogels that are ultimately wasted, inhibiting the cell densities attainable. Furthermore, in common with other bioprinting technologies, this approach has thus far been unable to assemble microscale layered tissue using collagen, the primary human Extra Cellular Matrix (ECM) material. An innovative alternative method is centrifugal casting, where hydrogels are applied to the inside of a closed cylinder that is then rotated. Centrifugal forces push the hydrogel to coat the internal surface creating a tube, however layer thicknesses were limited to 1mm as dictated by the volume added. The technology has received limited attention since the initial pioneering work by Mironov *et al*.^19,23,24^. Centrifugation has also been more recently exploited as part of the sacrificial writing into functional tissue (SWIFT) technology to create high-density macroscale constructs with microscale channels^25^.

Building on these past examples, we present a novel biofabrication technology termed rotational internal flow layer engineering (RIFLE) that enables the rapid, high-resolution bioassembly of tuneable, microscale hydrogel and high-density cell layers into a macroscale tubular construct. By exploiting the fluid dynamic state of rimming flow, cell-laden hydrogels were suspended as microscale liquid layers on the internal surface of a rotating tube. The addition of a gelation process, such as an ionic crosslinking reagent or temperature change, gels the layer before the addition of further layers. Progressively, a multi-layered tubular structure is built up, mimicking the macro and microscale architectures of native tissues.

## Results and Discussion

### Development of the RIFLE process

The equipment and process are shown schematically in (Fig. 1A) and in the animation in Supplementary Movie 1. Initial process development was carried out using alginate hydrogel and calcium chloride crosslinker before moving onto vascular tunica media tissue bioassembly with collagen. The addition of liquid phase alginate and calcium chloride into rotating closed cylinders failed to produce multiple layers due to variable hydrogel volume addition forming inconsistent thicknesses and because there was no route for the excess gelation agent to escape. The design of the mould was therefore modified to include drain holes to allow excess fluid to flow out of the cylinder. Liquid remaining within the cylinder was suspended as a thin, stable film on the rotating surface in a fluid dynamic process known as rimming flow^26^. A stopper tube fitted into the end of the tubular moulding ensured that outflow of excess fluid was possible only through the drain holes and also constrained the gelled layers within the tube. A component and reagent list can be found in supplementary information, table S1.

**Fig. 1.**
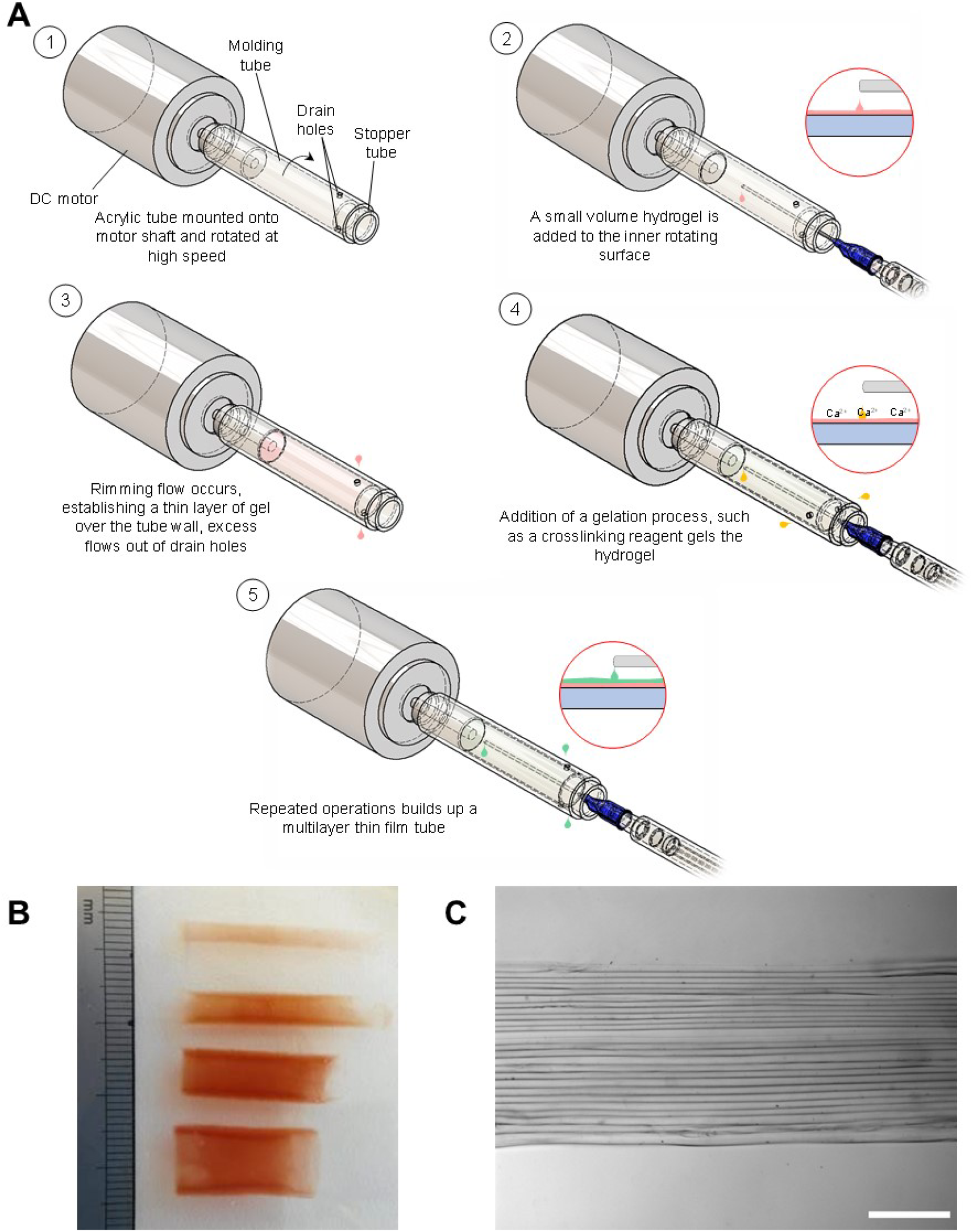
Illustration of RIFLE equipment and process. (**A**) Schematic representation of the RIFLE process. (**B**) Layered alginate tube size examples formed from moulds of different internal diameters, note: red food dye added to alginate to improve visualisation. (**C**) Brightfield microscopy image of a tube wall showing discrete alginate layers, white scale bar is 500μm.

Alginate hydrogel drops were observed landing on and then spreading over the inner rotating mould surface before excess liquid exited through the drain holes. A similar process was then observed for crosslinker calcium chloride, which gelled the previously applied alginate layer. Repeated alternate applications of alginate and calcium chloride built up a layered tube. The use of a range of moulds of various inner diameters allowed the assembly of example tubes of different diameters (Fig. 1B). Brightfield microscopy of tube walls revealed stratified microscale layers (Fig. 1C). A layered tube took less than 20 minutes to produce and required a low volume of hydrogel: a Ø10mm, 35-layer tube formed at 9000rpm used 1.5ml of 1% (w/v) alginate.

The observed rimming flow transition of the liquid from drops to thin layers on the internal rotating surface is most widely known in the industrial process of rotational moulding, where molten plastics are formed into large scale, single layer, hollow components^27^. Liquids rotated in a mould at high speeds will transition into rimming flow, suspending a uniform layer of liquid across the inner surface^28^. Here we have applied this principle to the formation of multi-layered hydrogel tubes, whilst allowing excess fluid a route to escape using drain holes. The fluid dynamics of rimming flow in closed cylinders has been studied extensively from a theoretical perspective^26,29,30^ and experimentally^31,32^. However, to our knowledge, no study has included the use of drain holes to allow the flow of excess liquid from the cylinder to establish thin layers or to use it as a method for forming microscale multi-layered structures.

### Rotational speed and layer thicknesses

Rotational speed is a key parameter for the industrial process of rotational moulding^27^ and in studies investigating the rimming flow behaviour of liquids in rotating closed cylinders^28,33^. We aimed to determine whether the rotational speed of the moulding tube influenced the thickness of the layers formed. Layered alginate tubes were made at speeds of 4500, 6000, 7500 and 9000rpm. Layer thicknesses were found to be a function of the speed of rotation: microscopy images of layers in tubes formed at different speeds are shown in (Fig. 2). Analysis of the layer thicknesses revealed that increasing rotational speeds formed thinner layers (Fig. 2E).

**Fig. 2.**
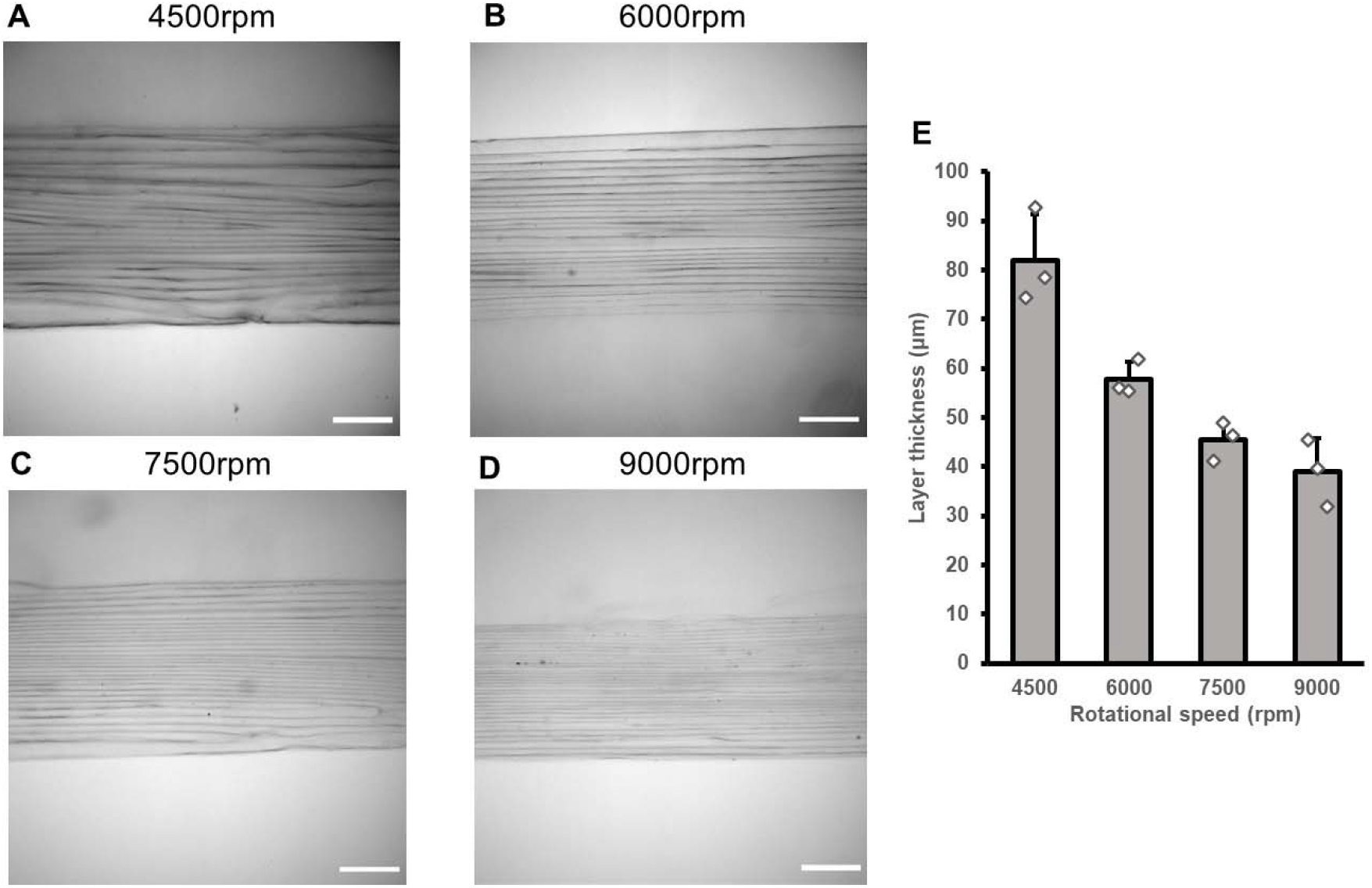
Layer thicknesses for RIFLE are determined by the speed of rotation. Brightfield microscopy images of tube walls formed using alginate 1% (W/V) at various rotational speeds. Scale bars are 500μm. (**A**) 4500rpm, (**B**) 6000rpm, (**C**) 7500rpm and (**D**) 9000rpm. (**E**) Plot of layer thicknesses for rotational speed variation. Mean with error bars ±SD n= 3.

As previously stated, no directly comparable theoretical or experimental fluid dynamic study is available. However, higher rotational speeds have been shown to reduce the thickness of layers for liquids adhered to the external surface of rods^20,21,28^ and also to promote the even spreading of molten plastics in rotationally moulded parts^34^. It is reasonable to assume similar fluid dynamic effects influence the thickness of the layers formed in multilayer constructs made using RIFLE. Rotational speed variation therefore provided a method of controlling layer thickness and thus the microscale dimensions of bioassembled stratified tissue. Such dimensional tunability is an important feature when attempting to mimic native stratified tissue with a range of layer thicknesses found in different examples.

### Alginate concentration and layer thicknesses

The concentration of liquid-phase alginate has been shown to influence multiple properties of bioprinted constructs, including elasticity, degradation and the behaviour of encapsulated cells^35^. Alginate concentrations of 4%, 2%, 1% and 0.5% (w/v) were used to form stratified tubular structures (Fig. 3). Analysis of the layer thicknesses revealed that the usage of higher percentage composition alginates resulted in tubes composed of thicker layers (Fig. 3E). Higher percentage concentration alginates are more viscous^36^, establishing a positive correlation with between layer thickness and viscosity. Higher viscosity materials have previously been shown to require greater speeds to make the transition to rimming flow in both closed cylinder experiments^37^ and theoretical models^28^. High viscosity alginates are likely to require greater centrifugal forces to form thinner layers.

**Fig. 3.**
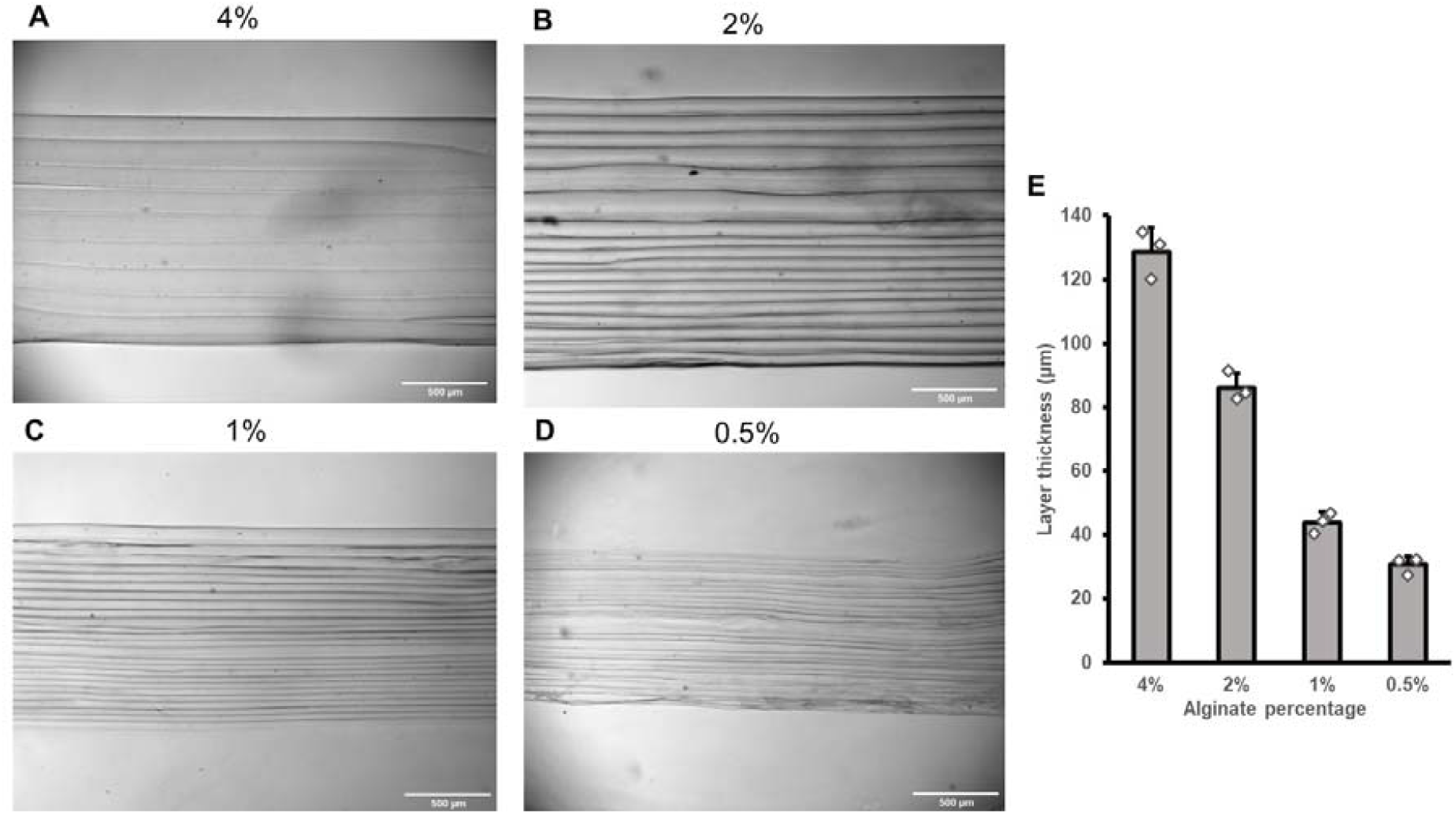
Alginate concentration influences the microscale layer thicknesses in tubes formed using RIFLE. Brightfield microscopy images of tubes walls with layer thicknesses and alginate concentration variation for tubes formed at 9000rpm. Alginate percentages (w/v) (**A**) 4%, (**B**) 2%, (**C**) 1% and 0.5%. Scale bars are 500μm. (**E**) Plot of layer thicknesses for alginate concentration variation. Mean with error bars ±SD, n = 3.

Hydrogel concentration is a key factor when considering the degradation profile of the printed tissue in a physiological environment, with desired faster or slower rates determined by the specific study or therapeutic objectives^38^. However, when seeking to use low viscosity hydrogels, such as low concentration alginate, collagen or decellularised extracellular matrix bio-inks^39,40^, researchers often find that they are unable to form microscale features using traditional bioprinting methods. A compromise between the desired hydrogel properties for tissue functionality and printability for the biofabrication technology is often needed^35,36^. The presented system is able to form the microscale feature of layers using low viscosity alginate by suspending the liquid phase in a stable layer prior to addition of a gelation agent.

### Encapsulated cell viability

Following development of the stratification process we sought to determine if cells could be included in microscale layers. HEK cells encapsulated into 1% (w/v) alginate layers were shown to have a viability of 97.5%±0.4 24 hours after RIFLE formation. The proportion of viable cells decreased in subsequent days to 60.4% ±1.2 at day 10 (Fig. 4E). The reduction in viabilities may be attributed to the formation of necrotic cores as cells proliferated and transitioned into larger spheroid bodies^41^ and has been previously observed in encapsulation protocols using alginate^22,42^.

**Fig. 4.**
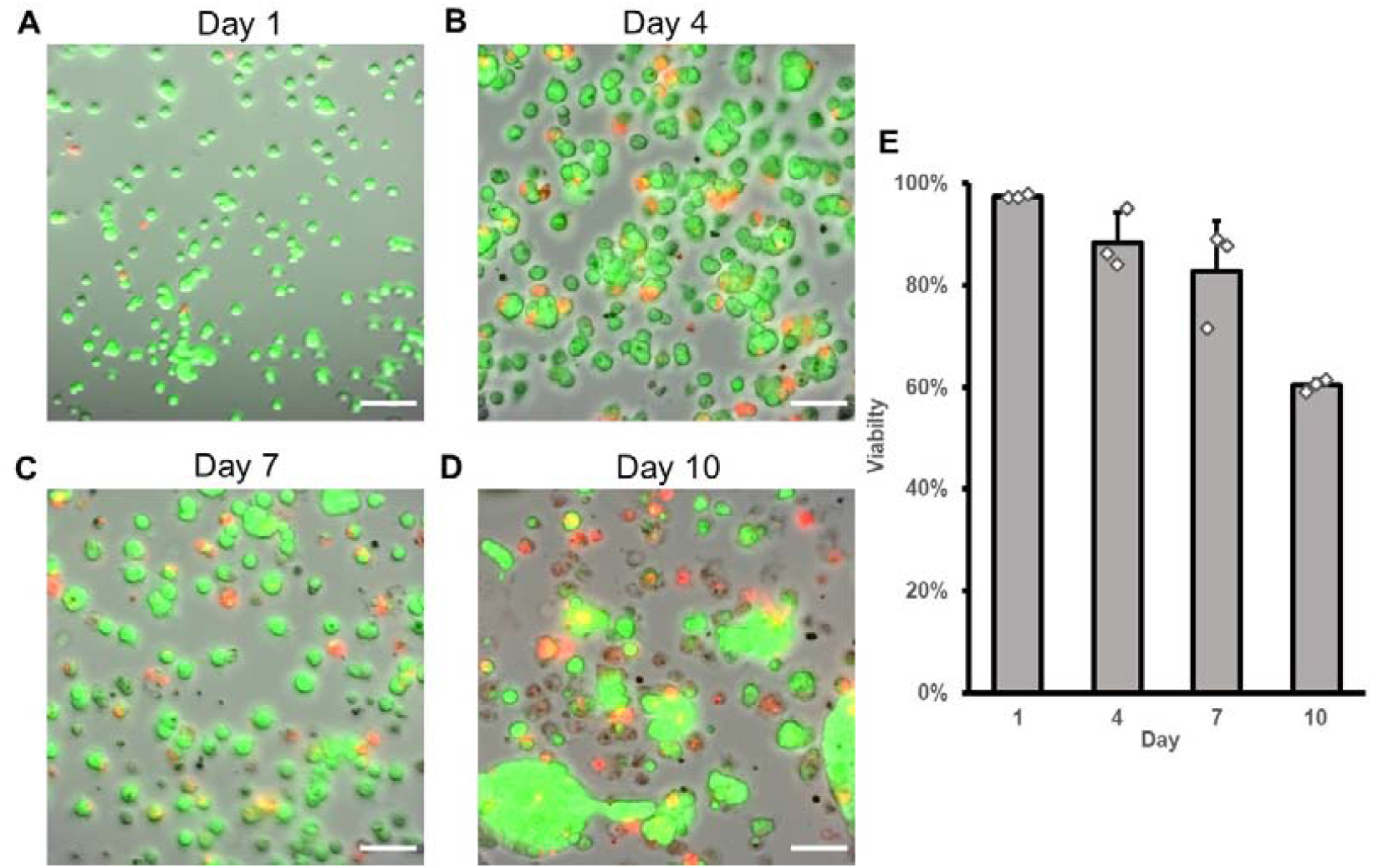
HEK cell viabilities following encapsulation into layers using RIFLE. Combined brightfield and fluorescent microscopy images showing HEK cell in tubes formed at 9000rpm, 1% alginate (W/V). Live (green) and dead (red) cells at (**A**) day 1, (**B**) day 4, (**C**) day 7 and (**D**) day 10. Scale bars are 100μm. (**E**) Plot of cell viability data following layer encapsulation. Mean with error bars ±SD, n = 3.

The data indicate that the overall forming protocol of trypsinisation, pelletisation, suspension in alginate and the RIFLE process results in a high cell viability with similar percentages to those attained using extrusion bioprinting, the most common biofabrication technology ^43,44^. Other cylindrical fabrication methods have presented viabilities in the first 24 hours after formation of 80% for rod dipping^22^, 95% for closed cylinder centrifugal casting^24^ and 20% to 60% for electrospinning^45^. Throughout the RIFLE process cells are subjected to centrifugal forces arising from tube rotation and also shear forces from fluid flow. Simple centrifugal force calculations reveal that a single cell is subjected to forces an order of magnitude lower than standard centrifugation used during pelletisation (1.55 × 10^−11^N vs 1.22 × 10^−10^N). Shear forces are also believed to have a detrimental impact on cell viability with Mironov et al. proposing that shear forces arising from cell transfer rather than centrifugal forces were responsible for cell death in their closed cylinder system^24^. The viability data demonstrates that shear forces are low enough for cell survival and indicates that cell encapsulation at higher rotational speeds than those presented here may be feasible in the future.

### Encapsulated cell layer positioning

Labelled high-density HEK cell populations - 10^8^ cells per ml – were assembled in varying patterns of 7 rainbow colours (Fig. 5A), alternating red and green single (Fig. 5B), double (Fig. 5C), triple (Fig. 5D) and quadruple (Fig. 5E) banded layers. Confocal microscopy in the sagittal plane revealed patterned layers of, red, green and rainbow colours cell populations, as assembled. Labelled red and green cells were also assembled in tubes with 10 and 5 acellular layers between them. Combined fluorescent and brightfield microscopy of the bioassembled tubes showed encapsulated cells populations distanced by 10 (Fig. 5F) and 5 (Fig. 5G) distinct layers, also as assembled. The results confirm that each cell layer is formed as an individual process and is discrete from the surrounding adjacent layers, allowing the formation of heterogenous composite layered structures. High-density cell layers were observed to be thicker than acellular and low-density layers. Previous studies have highlighted a positive correlation between cell density and hydrogel viscosity^46^, with the effect here that the RIFLE process forms thicker layers. The low volume of hydrogel needed for individual layer assembly enables high density cell encapsulation to be attained. The presented density of 10^8^ cells/ml aligns with observations in native cardiac tissue^47^ and is at least an order of magnitude higher than previous tubular biofabrication technologies that require large volumes for emersion, spraying or extrusion^20–22,48^.

**Fig. 5.**
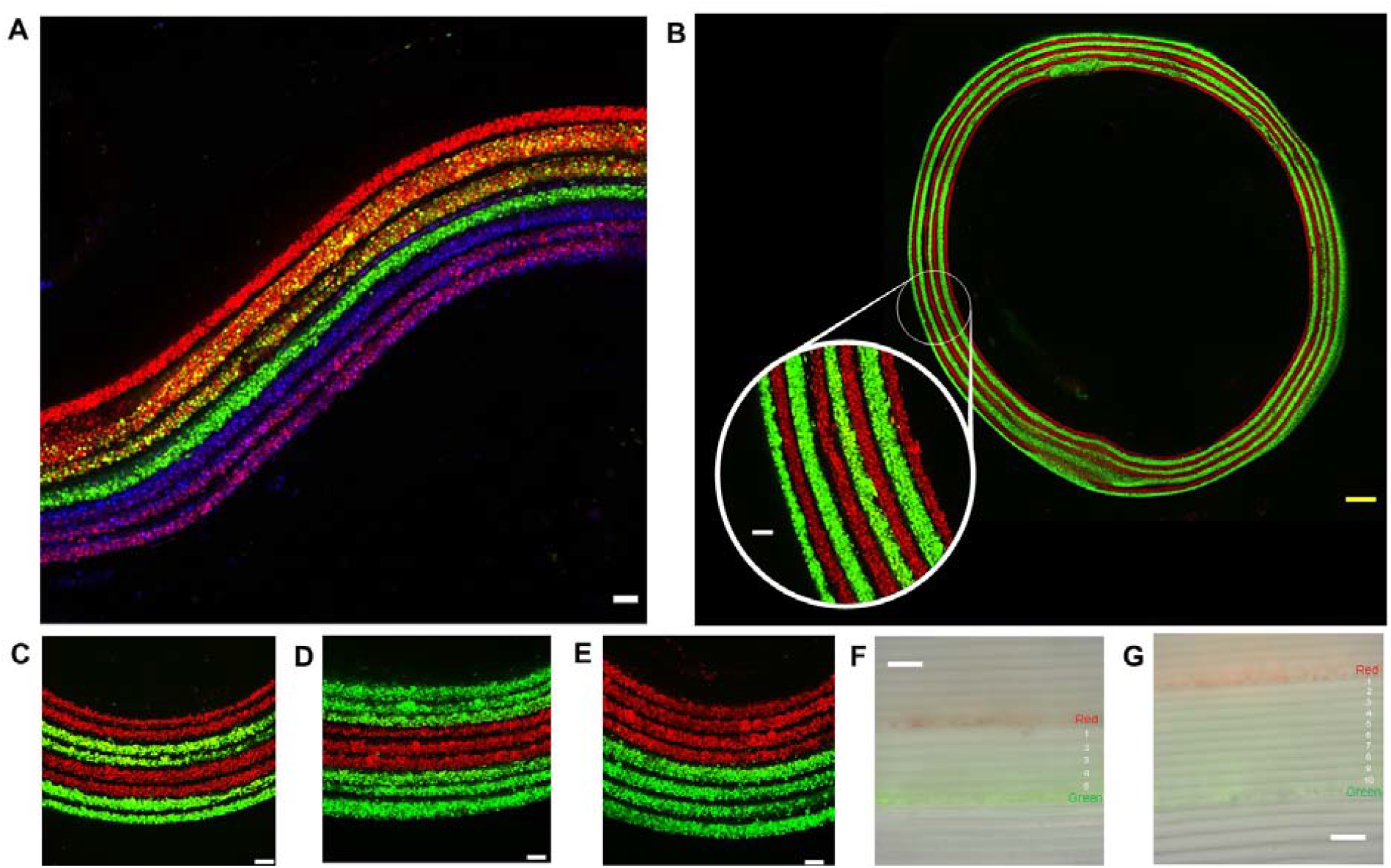
Cell layer positioning using RIFLE. (**A**), (**B**), (**C**), (**D**) and (**E**) Sagittal plane confocal images of patterned layers made using prelabelled red, green and blue high density HEK cells (10^8^ cells/ml) in 1% alginate (w/v) tubes. (**F** and **G**) Combined brightfield and fluorescent microscopy images in the transverse plane showing the position of red and green cell layers relative to one another with 5 or 10 acellular layers between as indicated. White scale bars are 100μm, yellow scale bar is 500μm.

Stratified tissue varies considerably throughout mammalian anatomy, with multiple combinational cell types, matrices and microlayer anatomy thicknesses. The data shows the bioassembly of high-density cells into microscale heterogenous structures, highlighting the potential to form a range of representative stratified 3D tissue. The precise spatial distancing of independent cell populations from one another may also have applications beyond tissue engineering, such as in the investigation of cell-to-cell signalling.

### Vascular tunica media tissue engineering with RIFLE

We sought to obtain data to determine if RIFLE was able to form layered vascular tubular structures using the native hydrogel collagen and encapsulated human smooth muscle cells (hSMCS). The intention was to mimic the native tunica media layer of vascular tissue, which is itself subdivided into further concentric microscale layers of hSMCs, known as the medial lamellar unit^49^. The same equipment for alginate bioassembly was used but with the process carried out in a 37°C environment. A 3 minute gelation time was applied for each collagen layer rather than the ionic crosslinker addition for alginate. Liquid neutralised collagen was observed spreading over the inner rotating surface of the tube with excess exiting from the drain holes in the same fashion as alginate. The macroarchitecture of completed collagen tubes was a tubular shape with an opaque white appearance (Fig. 6A). Acellular tubes formed showed evidence of microscale layer formation when viewed using brightfield microscopy. However, visualisation of layers was more difficult in comparison to alginate tubes due to the opacity of the gelled collagen, with layers of ≈15μm visible on the outer edge of the wall (Fig. 6B). The requirement to allow each layer 3 minutes to gel extends the bioassembly time, with 25 collagen layers taking approximately 1.5 hours.

**Fig. 6.**
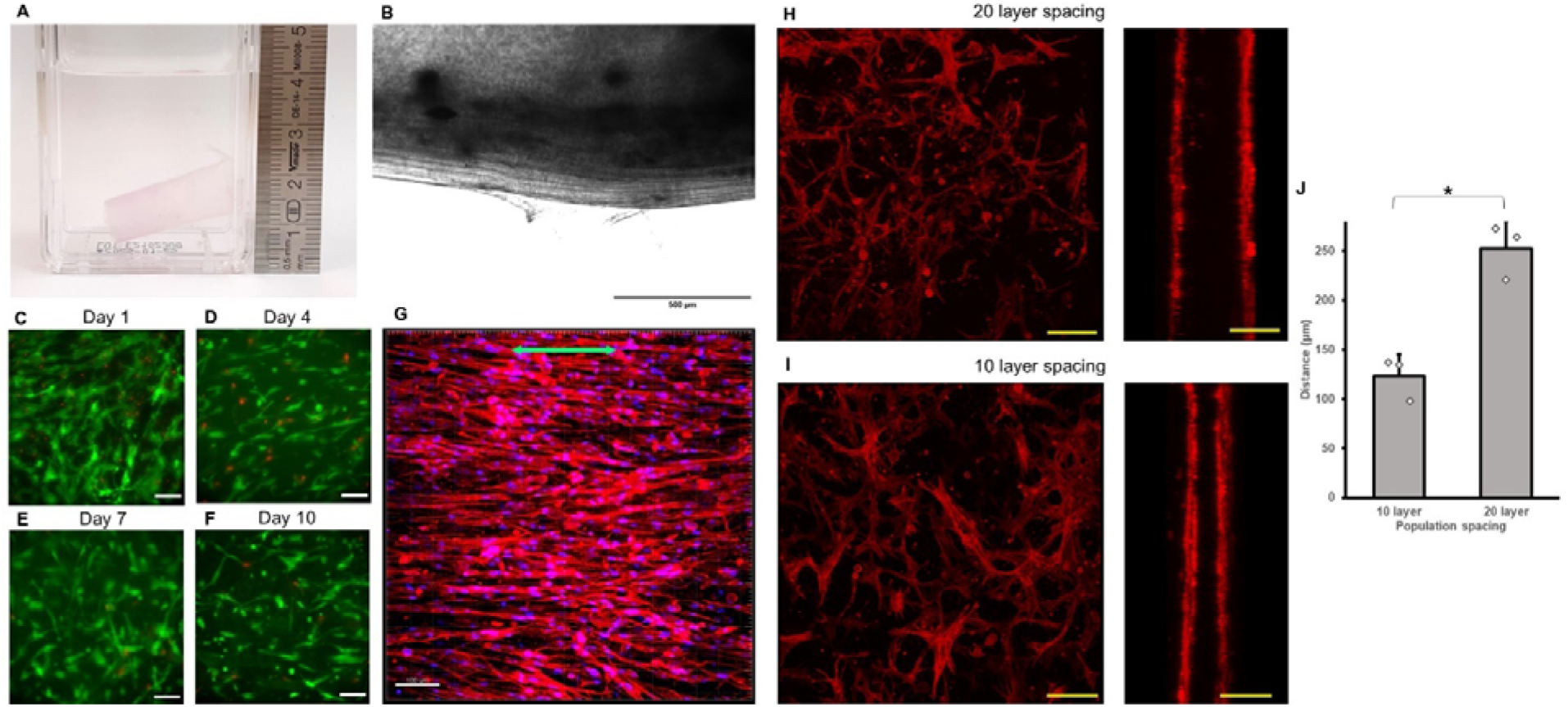
Bioassembly of vascular tunica media tissue using RIFLE. (**A**) Macroscale collagen tube architecture with an opaque appearance. (**B**) Light microscopy of microscale acellular collagen layer formation. Black scale bar is 500μm (**C** to **F**) Single layer hSMC viability in collagen. Live (green) and dead (red) cells at 1, 4, 7 and 10 days post assembly. White scale bars are 100μm (**G**) hSMC orientation 4 days after stratification. F-actin (phalloidin, red) nuclei (Hoechst, blue). Green arrow indicates circumferential direction. White scale bar is 100μm (**H** and **I**) Confocal stacks in the sagittal and transverse planes showing F-actin stained (phalloidin, red) hSMCs layers positioned with 20 and 10 acellular layer spacing. Yellow scale bars are 200μm. (**J**) Plot of measured distances between hSMC layers for the 10 and 20 layer groups. Mean with error bars ±SD, n = 3, two-sided student t test, p =0.0032. *P<0.005.

To bioassemble the vascular tunica media, hSMCs were suspended in neutralised collagen prior to addition to the inner surface of the rotating mould. hSMCs added as a single layer amongst acellular layers demonstrated viability for up to 10 days post assembly (Figs. 6C to 6F). Timelapse videos of hSMCs in the first 24 hours after bioassembly showed the cells transitioning from a spherical to a spread morphology in the plane of the layer (Supplementary Movie 2). Macroscale tubes containing hSMCs encapsulated in all layers were allowed to compact onto an inner mandrel for 4 days after bioassembly to prevent over shrinkage of the tube as viable cells haul up surrounding collagen fibrils^50^. Upon release from the mandrel F-actin visualisation revealed a network of hSMCs aligned in the circumferential direction (Fig. 6G). Such cellular orientation closely mimics the microscale organisation observed in native vascular tissue^15^. Alignment of cells in collagen has previously been shown to be influenced by mechanical stress^51^ and here the static tension induced by compaction onto the inner mandrel is likely to be contributing to the alignment of hSMCs.

To determine the thickness of assembled collagen layers, independent populations of hSMCs were assembled with 10 or 20 acellular layers spaced between them. F-actin visualisation via phalloidin staining and confocal microscopy revealed spread hSMCs in two distanced, independent, narrow cell-width microscale layers (Figs. 6H and 6I and Supplementary Movies 3 and 4). The distance between cell layers in the 10 and 20 layer assemblies increased proportionately (Fig. 6J), with an average layer thickness of 12.5±1.7 μm. This distance closely matches published measurements of the thickness of the native medial lamellar unit, recorded at 13.9±1.2 μm (rat)^15^ and 13.2μm (human)^49^. The assembly of viable hSMCs into concentric, anatomically accurate layers represents an important progression for vascular biofabrication technologies. Bioprinting studies frequently only refer to the macrolayers of the tunica intima, media or adventitia, oversimplifying the complex microtissue layered architecture within these structures, such as the medial lamellar unit^15,49,52^. The results show that RIFLE is able encapsulate vascular cells in anatomically accurate concentric medial lamellar units in the native ECM material of collagen.

There remain several limitations of the assembled tunica media tissue in comparison to native tissue. The constructs do not feature the elastin lamellar that lie between the medial lamellar units^15^. These sheets are believed to significantly contribute to the mechanical strength of vessel walls^53^. In addition, we have only tested a simple tubular shape whereas native macroscale tissue can be more complex, such as vascular bifurcations and ventricular bodies. The industrial process of rotational moulding utilises rimming flow in hollow moulds with complex geometries. Further investigation will be required to determine if more complex multi-layered macroscale geometry can be created using the RIFLE principle of rimming flow.

The data provides evidence that the RIFLE system can be directed towards vascular tissue engineering via the thermal gelation of microscale collagen layers into the medial lamellar unit. Microstratification of cells encapsulated in collagen is also significant for other tissue engineering applications, being the principal component of most extracellular matrices and accounts for 30% of human protein mass^54^. Furthermore, collagen has typically proved difficult to manipulate using existing bioassembly technologies^55^. This is due to a combination of low viscosity and extended thermal gelation time^13,16^, ultimately requiring a high level of self-support and temperature control when transitioning from a liquid to a gel^56,57^ a challenge prevalent throughout biofabrication, with technologies often requiring specific bioink hydrogel formulations. Collagen itself is often combined with other hydrogels such as alginate^58^ to enable bioprinting. The RIFLE system circumvents this problem for stratified tissue with the stable suspension of the liquid phase hydrogel in the desired microscale spatial constraint prior to application of the gelation reagent or process.

### Economical stratified tissue bioassembly

The described system uses simple equipment needing only the single axis rotation of a DC motor and therefore requires no complex control software or associated programming skills. The apparatus used is all readily available to research laboratories at a relatively low cost. The combined total for the equipment presented is less than $950, the majority of which is for the DC motor and controller, with cheaper versions available. The high cost of bioprinting equipment alongside required specialised expertise^59^ is often a barrier to researchers seeking to assemble tissue in the laboratory^60^ with current commercial options costing between $10,000 and $1,000,000 ^61^. It is hoped that more affordable, user-friendly technologies will enable greater access for laboratories wishing to develop engineered tissue for research and ultimately clinical purposes. The speed of layered tissue fabrication using RIFLE presents a further advantage to researchers too, especially in comparison to technologies such as cell-sheet engineering which requires extensive 2D culture^48^ and maturation times^5^. The described system therefore further supplements the number of emerging biofabrication technologies available to researchers seeking to mimic microscale tissue features. In this regard RIFLE can be considered to be an enabling technology for the two-step biomanufacturing roadmap outlined by Wolf et al^62^. Firstly, the creation of the tissue functional unit microarchitecture, the medial lamellar unit in the presented example, followed by the biosassembly of these units into a bulk tissue, here a macroscale tube. It is hoped that the future use of RIFLE will facilitate others to repeat this approach for a range of different stratified tissue types.

## Materials and Methods

### Equipment and process

Multi-layered tubes were fabricated by using a 24V DC motor (EC-i 30, Maxon Motor Ltd, UK) to rotate a clear acrylic tubular mould at high speeds. Motor speed was controlled using a DEC Module 24/2 (Maxon Motor), an Arduino microcontroller and a LabView (National Instruments, USA) program bespoke to the controller. Rotational speed in revolutions per minute (rpm) was determined through analysis of the digital outputs from the motor hall effect sensors using the same bespoke LabView program. With rotation set at the desired constant speed, liquid phase alginate (Protanal LF10/60FT, FMC Biopolymer, UK) was dropped onto the inner surface of the rotating mould using a 21ga blunt-end needle (Fisnar, UK). The internal diameter of the rotating mould was 10mm unless otherwise stated, with two 1mm diameter drain holes located opposite one another. Rimming flow evenly distributed the fluid over the surface of the rotating mould, establishing a suspended thin layer of hydrogel, with excess fluid flowing out of the drain holes. 100mM CaCl_2_ solution in H_2_O (BDH chemicals, UK) was then added to the mould in a similar fashion, gelling the suspended alginate layer, with excess also flowing out of the drain holes. Repeated alginate and CaCl_2_ addition sequentially built up a multi-layered tube. A smaller diameter stopper tube retained gelled layers within the mould. On completion the mould was removed from the motor and light tapping was sufficient to release tubes through the open end. Computer-aided design and animations were created using Autodesk Inventor Professional 2023 (Autodesk Inc, USA).

### Rotational speed variation

Acellular tubes were fabricated using 1% (w/v) alginate, at rotational speeds of 4500, 6000, 7500 and 9000rpm, with 3 independent replicates being made for each speed group. Images of each tube were captured with brightfield microscopy with a x4 objective using an Eclipse Ti2 microscope (Nikon Instruments Europe, Netherlands). Layer thicknesses were determined using ImageJ software by assigning a 100μm wide selection box across the wall image and running the “plot profile” function. Graphical visualisation of the profile enabled the relative positions of the layer boundaries to be discerned and individual layer thicknesses to be calculated and subsequently averaged.

### Alginate concentration

Acellular tubes were fabricated at 9000rpm using alginate concentrations of 0.5%, 1%, 2%, and 4% (w/v), with 3 independent replicates being made for each concentration group. Images of each tube were captured using brightfield microscopy with a x5 objective on a Zeiss Axiovert microscope (Zeiss, Germany). Layer thicknesses were determined as previously described.

### Cell culture

Human embryonic kidney 293T (HEK) cells were cultured in Dulbecco’s Modified Eagle Medium (Gibco, 41966-029) supplemented with 10% fetal bovine serum (Gibco, 10270-106) and 1% Penicillin Streptomycin (Gibco, 15070-063). Human Aortic Smooth Muscle Cells (354-05A, Cell Applications, San Diego, USA) were cultured in Smooth Muscle Cell Growth Medium (311-500, Cell Applications) and used at passages below 5. All cells were cultured at 5% CO_2_ and 37°C.

### Cell viability in alginate

Multi-layered tubes were formed with 1% alginate in PBS (Gibco 14190) at 9000rpm. HEK cells were suspended in alginate at a density of 2.8 × 10^6^ cells/ml and were included as a single central layer between acellular layers. Completed tubes were incubated in cell culture media at 5% CO_2_ and 37°C. Sections were removed from 3 separate tubes (n of 3) at 1, 4, 7 and 10 days after formation and a Live/Dead assay (Abcam, UK, ab115347) was performed according to manufacturer’s instructions. Images were captured using an Eclipse Ti2 microscope (Nikon) with a x20 objective. Viability was calculated as the ratio of the area of live and dead cells determined through the application of the ImageJ software “make binary” and “analyse particles” functions to the green (live) and red (dead) channel images^63^.

### Cell layer positioning

Flasks of HEK293 cells were pre-labelled red or green using cell tracker labels (C7025 and C34552, Thermofisher Scientific) or blue using Hoescht 33342 (1:1000, H3570, Invitrogen, UK) according to manufacturer’s instructions. Labelled cells were suspended in 1% (w/v) alginate in PBS and included in 9000rpm bioassembled tubes. For adjacent patterned high-density cell layers HEK cell populations at a density of 1.0 × 10^8^ cells/ml were included as a central band in varying patterns between acellular outer and inner layers. For rainbow patterns, red, green and blue labelled HEK293 cells in 1% (w/v) alginate were mixed in ratios matching the RGB values for red (100, 0, 0) orange (66, 34, 0), yellow (50, 50, 0), green (0, 100, 0), blue (0, 0, 100), indigo (34, 0, 66), and violet (41, 0, 59) to create 7 independent cell colour populations^64^. Tubes were sectioned and imaged in the sagittal plane using a Nikon A1R confocal inverted microscope in galvano scan mode with a x4 or 10x objectives and a x4 tile scan mode for whole ring imaging. For spaced layer positioning HEK293 cells at a density of 2.7 × 10^6^ cells/ml were layered between acellular layers with 10 or with 5 acellular layers between an outer green and an inner red population. Images were captured after 24 hours of culture using an Eclipse Ti2 microscope (Nikon) using a x4 or x20 objective.

### Collagen layer tube formation

The motor and rotational mould assembly were placed into a humidified chamber set at 37°C. Collagen type I was prepared in small batches, transferred to a syringe on ice and used immediately to prevent premature gelation. 445μl Rat Tail collagen I at 3mg/ml in 0.02N acetic acid (Corning, USA) was neutralised with a 55μl 2:1 ratio mixture of x10 DMEM (Sigma D2429) and 0.4M NaOH (Sigma S5881) and then 0.4M NaOH dropwise until the phenol red in the DMEM indicated a neutral pH had been reached. Liquid phase collagen was dropped onto the inner rotating surface of the 4500rpm rotating mould in the same fashion as alginate protocols and allowed to gel for 3 minutes before addition of the next layer. Microscopy images were captured using an Eclipse Ti2 microscope (Nikon) using a x4 objective.

### Tunica media tissue assembly

For viability studies in collagen, hSMCs were suspended in collagen at a density of 7.4 × 10^6^ cells/ml and included as a single central layer between acellular layers. Completed tubes were incubated in cell culture media at 5% CO_2_ and 37°C. Sections were removed from 3 separate tubes (n of 3) at 1, 4, 7 and 10 days after formation and a Live/Dead assay (Abcam, UK, ab115347) was performed according to manufacturer’s instructions. Images were captured using an Eclipse Ti2 microscope (Nikon) with a x20 objective. The high difference in morphology and density between live, (spread, large and overlapping) and dead cells (rounded, small) prevented accurate quantification and viability was therefore only qualitatively analysed. For timelapse imaging hSMCs were included in a single layer at a density 1 × 10^6^ cells/ml and brightfield images taken at 5-minute intervals for 24 hours immediately after bioassembly using a x20 objective and an Eclipse Ti2 microscope (Nikon) with a stage top incubator. For cell orientation imaging hSMCs at a density of 0.6 × 10^6^ were formed into 22 collagen layers and the tube placed over a rigid Ø8.5mm mandrel to prevent over compaction. After 4 days the construct was removed from media, washed with PBS and fixed with 4% paraformaldehyde for 15 minutes at room temperature, before permeabilisation with 2% triton for 15 minutes at room temperature. Staining of actin filaments was achieved using Phalloidin-iFluor 594 (1:1000, ab176757, Abcam), nucelli were stained with Hoescht 33342 (1:1000, H3570, Invitrogen, UK). Images were captured using a Nikon A1R confocal inverted microscope in galvano scan mode with a 10x objective and visualised using IMARIS software (Bitplane, Switzerland). For medial lamellar unit layer width determination, independent hSMC populations at a density of 1.5 to 2 × 10^6^ cells/ml were included in collagen tubes with 10 or 20 layers spaced between them, 3 replicate tubes per spacing condition. After 24 hours the constructs were fixed, stained with phalloidin and imaged as detailed previously. Confocal stacks were visualised and the distance between layers ascertained using the imageJ 3D project function to rotate stacks through 90° to display the two layers in profile. The plot profile function was then applied to display the x positions of the peak fluorescent intensities from the two layers and the values subtracted from one another to calculate the total distance between layers.

### Statistical analysis

All data are shown as the mean with ±Standard Deviation (SD). Statistical analysis was performed using a student’s two tail t test.

## Supporting information

Supplementary Movie 1 - Animation of RIFLE equipment and process

Supplementary Movie 2 - 24 Hour post bioassembly timelapse of hSMCs in a single layer

Supplementary Movie 3 - Medial lamellar units - 10 layer spacing

Supplementary Movie 4 - Medial lamellar units - 20 layer spacing

## Data availability

Data and 3D models will be made publicly available on, an online data repository managed by the University of Edinburgh.

## Acknowledgements

We would like to thank Anisha Kubasik-Thayil at the University of Edinburgh IMPACT Imaging Facility where confocal imaging was carried out.

## Funding

We would like to acknowledge the following funding. I.H and J.D European Research Council (ERC) under the European Union’s Horizon 2020 research and innovation scheme (grant agreement no. 801041, Cyber Genetic Tissue Engineering), I.H and W.S UK Engineering & Physical Sciences Research Council (EPSRC) Doctoral Prize Fellowship (Grant No. EP/N509760/1) I.H and J.D UK Biotechnology & Biological Science Research Council (BBSRC) (grant agreement no.

BB/M018040/1).

## Author contributions

I.H, W.S and J.D designed the research. IH performed the experiments, analysed the data and wrote the manuscript with input from W.S and JD.

