## Supplementary Materials: Table S1 for "Stratified tissue biofabrication by rotational internal flow layer engineering"

Table S1. Component and reagent list for RIFLE.

| **RIFLE components** | **Supplier** | **Part No.** |  |
| --- | --- | --- | --- |
| 30V DC EC-I Maxon motor | Maxon Motor Ltd | 539473 |  |
| Maxon, DC Motor Controller | Maxon Motor Ltd | 367661 |  |
| Arduino, Mega 2560 | RS components | 7154084 |  |
| 0-30V, 0-5A, DC power supply | RS components | 1757368 |  |
| Clear acrylic tube OD13mm ID10mm | RS components | 2822256 |  |
| Clear acrylic tube OD10mm ID7mm | RS components | 2822234 |  |
| 21 gauge 1.5" blunt end needle | Fisnar | 040108000126 |  |
| 1ml luer lock syringe | Fisher Scientific | 15489199 |  |
| **Reagents** | **Supplier** | **Part No.** | **Lot Number(s)** |
| Sodium Alginate Protanal | FMC Biopolymer | LF10/60FT |  |
| Calcium Chloride | BDH Chemicals | 27587 | 011490K |
| Dulbecco’s Modified Eagle Medium | Gibco | 41966-029 | 2340247 |
| Fetal Bovine Serum | Gibco | 10270-106 | 2232250 |
| Penicillin Streptomyocin | Gibco | 15070-063 | 02096F18 |
| Smooth Muscle Cell media | Cell Applications | 311-500 | 22404, 22303 |
| Trypsin - EDTA solution | Gibco | 15400-054 | 2457694 |
| Phosphate Buffer Solution | Gibco | 14190169 | 2477761, 2526660 |
| Live and Dead Cell Assay | Abcam | ab115347 | 2101004218 |
| Cell tracker - Green | Thermofisher | C7025 | 2110523 |
| Cell tracker - Red | Thermofisher | C34552 | 1370420 |
| Collagen Type 1 | Corning | 354236 | 1308003,1147001 |
| x10 DMEM | Sigma | D2429 | RNBJ4442 |
| Sodium Hydroxide | Sigma | S5881 | 014K0006 |
| Phalloidin | Abcam | ab176757 | GR3406338-7 |
| Hoescht | Invitrogen | H3570 | 1932420 |
| Triton 100X | Sigma | T-8787 | 64H2616 |
